# Branched-chain ketoacid antioxidants mediate disease tolerance to sepsis

**DOI:** 10.1101/2025.05.22.654748

**Authors:** Aditya Misra, Efi M. Weitman, Michal Weitman, Jeannette R. Brook, Joseph M. Rone, Boryana Petrova, Randall T. Mertens, Narae Hwang, Katherin Zambrano Vera, Mark A. Perrella, Rebecca M. Baron, Naama Kanarek, Roni Nowarski

**Author notes:** These authors contributed equally (Aditya Misra, Efi M. Weitman, Michal Weitman).

## Abstract

Metabolic adaptation is crucial for surviving systemic infection and withstanding the pathological host response to infection known as sepsis. The liver orchestrates key metabolic adaptation programs that enable disease tolerance in sepsis, yet the impact of liver metabolites on sepsis susceptibility is not well understood. By broadly profiling liver metabolite landscapes in mice surviving or succumbing to bacterial sepsis, we found that dysregulation of the branched-chain amino acid metabolite family is associated with sepsis non-survival. Administration of branched-chain ketoacids (BCKAs) during *Klebsiella pneumoniae*-induced sepsis in mice enhanced survival, yet not through enhanced bacterial clearance or BCKA catabolism. Instead, BCKAs served as antioxidants by directly neutralizing hydrogen peroxide, preventing tissue lipid peroxidation. Targeted metabolomics in sepsis patients revealed low BCKA abundance as an early prognostic biomarker of sepsis non-survival. Our results identify BCKAs as a systemic shield against oxidative damage and highlight new metabolite targets to enhance disease tolerance to sepsis.

## INTRODUCTION

Sepsis is a dysregulated host response to infection that leads to organ failure and ∼20% of global deaths^1–3^. The major challenge faced by the host is how to effectively eradicate the invading pathogen while preventing detrimental damage to organs. Two adaptation programs are therefore set off during sepsis, one that mobilizes the immune system to neutralize the pathogen (‘resistance’), and one that limits the negative impact of infection on the host (‘disease tolerance’)^4^. Intriguingly, the clinical progression of sepsis is attributed mostly to inadequate host response to infection rather than the direct action of the invading pathogen^5,6^. This notion is underscored by highly overlapping molecular responses to a large diversity of pathogens, and the persistence of disease even in the absence of detectable pathogen^6–8^. Thus, the capacity to employ disease tolerance mechanisms during sepsis is an important defense feature crucial for survival.

Early after infection, the host prioritizes the elimination of the pathogen, which may come at the expense of disease tolerance and contribute to organ failure^8^. This conflict is rooted in the differential metabolic requirements for the two programs, where resistance mechanisms rely on anabolic biosynthetic pathways that consume energy, while disease tolerance mechanisms are dependent on catabolic macromolecule breakdown to produce energy^9^. Dysregulation of metabolic pathways during sepsis therefore compromises both immune-mediated pathogen clearance and tissue disease tolerance^8,10–12^. In order to maintain energy homeostasis during sepsis, the liver sustains glucose production via gluconeogenesis while limiting iron availability and toxicity ^11^. In parallel, hepatic PPARα activation mediates increase in mitochondrial fatty acid oxidation and a corresponding decline in glycolysis^13^, promoting protective ketogenesis while limiting the generation of proinflammatory metabolites and lactate that lead to tissue injury^14,15^. These liver metabolic adaptations are essential to establish disease tolerance to sepsis.

Alas, sepsis-induced metabolic dysregulation is an important driver of disease pathogenesis ^16,17^. Sepsis leads to metabolic exhaustion in various immune and other cells—a state in which cells are unable to maintain their energy demands ^8,10,12^. Additionally, systemic metabolism is shifted in sepsis, as reflected in the plasma metabolome, and can provide markers for sepsis severity ^18,19^. The shift in systemic metabolism may be partially attributed to sepsis-induced liver injury, which can predict mortality in sepsis patients^20–22^. Despite this compelling cellular and systemic evidence, the role of metabolites in driving or preventing the metabolic crisis during sepsis is underexplored. Metabolites are critically required to neutralize reactive oxygen species (ROS), that are among the most pathological systemic factors derived from local inflammatory environments^14,23,24^. ROS attack polyunsaturated fatty acids in cell or organelle membranes, as well as proteins and DNA, and have been strongly associated with sepsis mortality^25–28^. Hydrogen peroxide (H_2_O_2_) is a key antimicrobial effector molecule produced by immune and other cells to eliminate pathogens and is normally metabolized intracellularly by the glutathione redox cycle^29^. However, in sepsis, cellular glutathione pools are rapidly depleted by excess peroxide production^30,31^. This leads to high peroxide concentrations inside cells, causing cell death and peroxide release into the systemic circulation^32,33^. Loss of physiological oxidant control during sepsis is further driven by a decrease in the plasma antioxidant status, which particularly contributes to lipid peroxidation through the excessive activity of peroxides^25,34,35^. Despite the central role of ROS in sepsis pathogenesis, several clinical trials evaluating antioxidants as therapeutic agents showed low efficacy and high toxicity^36–38^. Thus, new antioxidant approaches that bypass cellular toxicity are urgently needed for sepsis therapy, particularly by enhancing physiological antioxidant mechanisms. Here, we describe the antioxidant and tissue protective role of branched-chain ketoacids, highlighting their potential use as biomarkers and therapeutic agents in sepsis.

## RESULTS

### Defining liver metabolite profiles associated with sepsis survival

Liver dysfunction is a common hallmark of sepsis, but how liver metabolites impact disease outcomes in sepsis is not well understood. Seeking to address this gap, we profiled liver metabolites in 43 age-matched male and female mice grouped based on their ability to control *Klebsiella pneumoniae* (*Kp*) sepsis (healthy survivors, S; terminally ill non-survivors, NS) (**Figure 1A**). Using ultrahigh performance liquid chromatography coupled to tandem mass spectrometry (UPLC-MS/MS), we profiled 917 metabolites across multiple metabolite families, including lipids, amino acids, nucleotides, xenobiotics, carbohydrates, cofactors and vitamins, peptides, and energy (**Figure 1B**). Principal component analysis (PCA) of the feature matrix showed clear separation between S and NS mice, implying that each group has distinct liver metabolic profiles (**Figure 1C**). Metabolite subgroups that were most highly enriched in NS mice were associated with fatty acids, fatty acid metabolism (acyl carnitine, dicarboxylate), and branched-chain amino acid (BCAA) metabolism. In contrast, fructose, mannose and galactose metabolism, bile acids, benzoate metabolism, and monoacylglycerols were enriched in S mice (**Figure 1D**). To further evaluate the important hepatic metabolite differences between S and NS mice, we built a multiple linear regression model using body temperature at the experimental endpoint as a continuous variable that serves as a proxy for disease severity. Univariate correlation showed that 7 of the top 20 metabolites (35%) that correlated with temperature belonged to the BCAA metabolism pathway (**Figure 1E**). We also implemented Metabolite Set Enrichment Analysis (MSEA) to characterize pathway-level changes between S and NS mice. MSEA similarly highlighted BCAA metabolism as a highly enriched pathway among differentially abundant metabolites (**Figure 1F**). These results suggest that metabolic products of the BCAAs leucine, valine, and isoleucine may either reflect or regulate sepsis disease severity.

**Figure 1.**
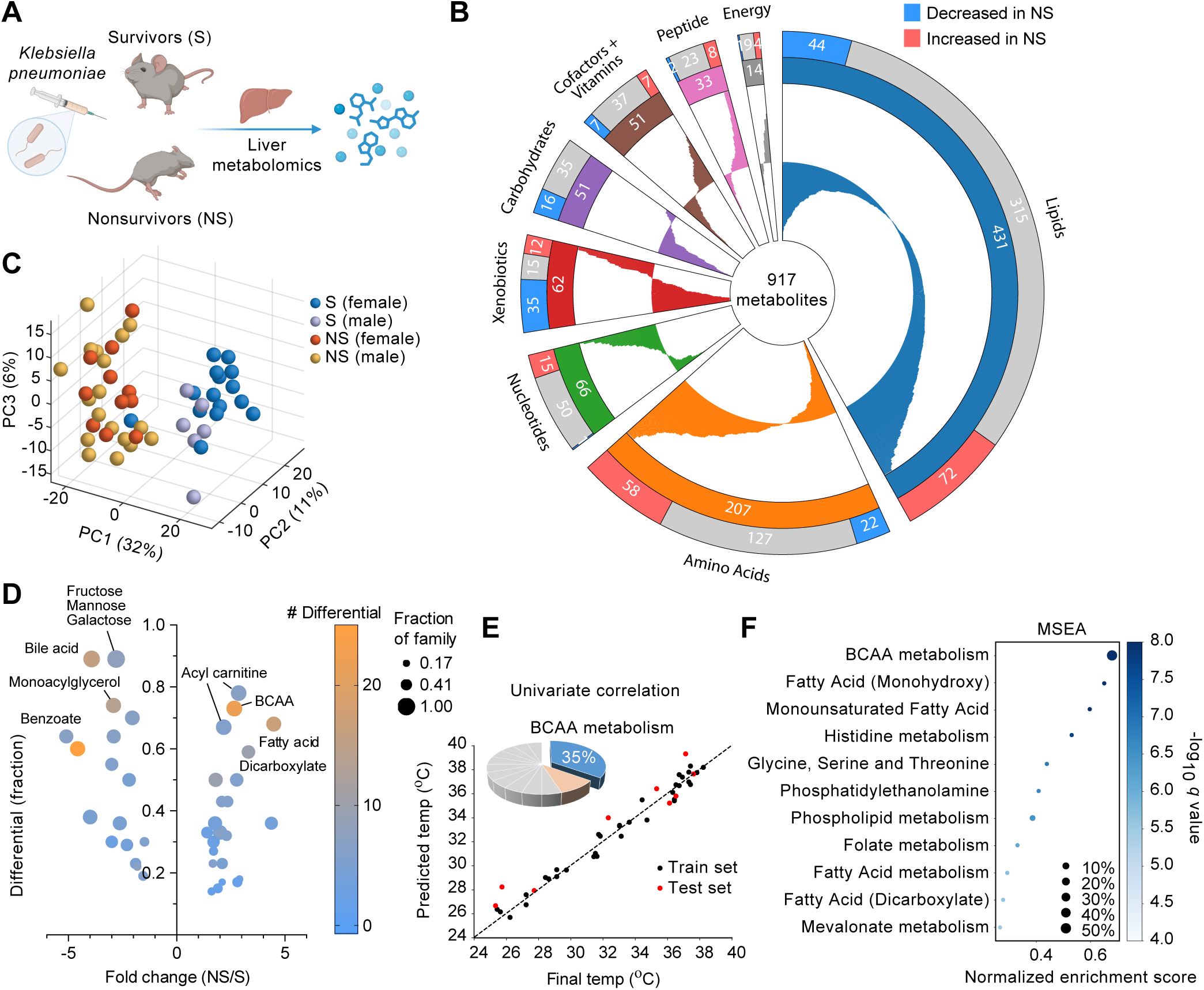
Defining liver metabolite profiles associated with sepsis survival. (A) Experimental scheme for targeted liver metabolomics in mice surviving (S) or not surviving (NS) *Klebsiella pneumoniae*-induced sepsis (*n*=43). (B) Relative abundance of 917 metabolites identified by ultrahigh performance liquid chromatography coupled to tandem mass spectrometry (UPLC-MS/MS) in livers of NS vs S mice across major metabolite families. (C) Principal component analysis (PCA) of metabolite distribution in S and NS mice. (D) Metabolite subgroup analysis according to differential metabolite abundance and the fraction of metabolites showing differential abundance comparing NS and S mice. (E) Univariate correlation between metabolite abundance and body temperature at end point. (F) Metabolite set enrichment analysis (MSEA) of metabolic sub-pathways differential in NS vs S mice.

### Branched-chain ketoacids rescue sepsis survival

BCAAs are mainly consumed through the diet and circulate in the blood. They can be stored in skeletal muscle where the enzyme branched-chain amino acid transaminase 2 (BCAT2, BCATm) converts BCAAs into the branched-chain ketoacids (BCKAs) 4-Methyl-2-oxopentanoic acid (2-Ketoisocaproic acid, KIC), 3-Methyl-2-oxopentanoic acid (2-Keto-3-methylvaleric acid, KMV), and 3-Methyl-2-oxobutanoic acid (2-Ketoisovaleric acid, KIV) (**Figure 2A**). During times of metabolic stress, the skeletal muscle breaks down and releases BCKAs into the bloodstream. Since the liver has high activity of the enzyme branched-chain ketoacid dehydrogenase complex (BCKDC), it has the capacity to attach CoA to the BCKAs and commits them to the TCA cycle as energy source (**Figure 2A**). To functionally assess the role of BCAA pathway metabolites in sepsis pathogenesis, we began by providing mice with daily intraperitoneal injections of a 1:1:1 equimolar cocktail of BCAAs or BCKAs during the course of *Kp*-induced sepsis. We found that BCKA supplementation protected mice from weight loss and death during the course of infection with *Kp* at lethal dose (LD)70 (**Figure 2B, C**). In contrast, BCAA supplementation did not provide a protective benefit over the PBS-treated control mice (**Figure 2B, C**). BCKA supplementation was also protective in mice infected with *Kp* at LD100, reaching an optimal effective dose of 0.25 mmol/kg or lower total BCKAs (**Figure 2D**).

**Figure 2.**
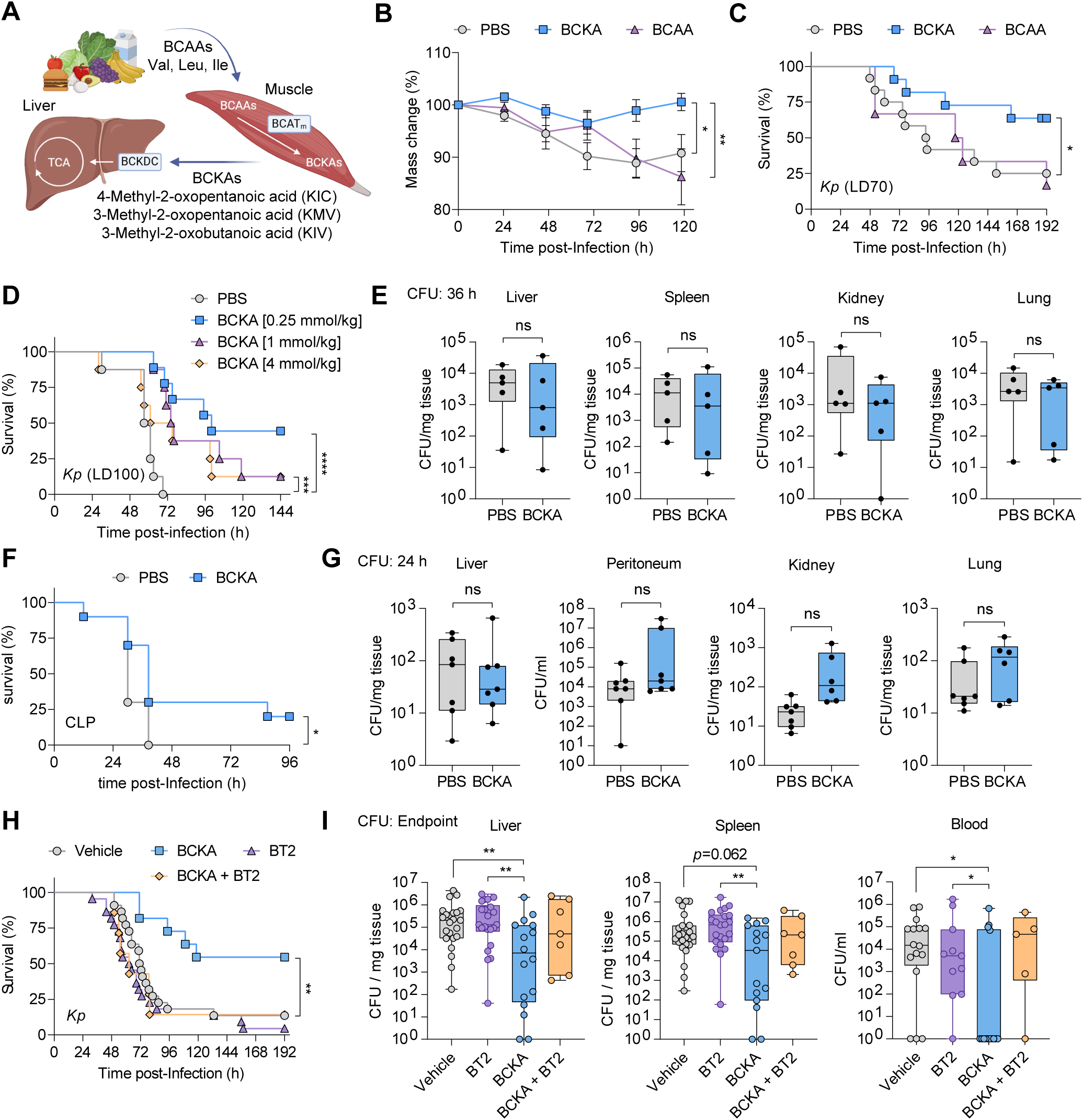
Branched-chain ketoacids rescue sepsis survival. (A) Scheme of branched-chain amino acid (BCAA) metabolism. Val, valine; Leu, leucin; Ile, isoleucine; BCATm, branched-chain amino acid transaminase 2 (BCAT2); BCKDC, branched chain ketoacid dehydrogenase complex. (B) Weight change of mice treated with 1:1:1 equimolar cocktail of BCAAs or BCKAs and infected with *Kp* at a lethal dose (LD)70 (*n*=7-12). (C) Survival of mice infected as in (B). (D) Survival of mice treated with the indicated BCKA doses and infected with *Kp* at LD100 (*n*=8-9). (E) Bacterial colony forming units (CFUs) in organs harvested 36 h post *Kp* infection. (F) Survival of mice induced with lethal cecal ligation and puncture (CLP) sepsis (*n*=10). (G) Bacterial CFUs in organs harvested 24 h post CLP induction. (H) Survival of mice treated with BCKAs and the branched-chain ketoacid dehydrogenase kinase (BCKDK) inhibitor BT2 (20 mg/kg) (*n*=7-22). Data shown as mean±SEM. *p<0.05; **p<0.01; ***p<0.001; ****p<0.0001.

BCKAs may improve survival by enhancing resistance (bactericidal) mechanisms or disease tolerance. To better understand BCKA protective function, we analyzed the bacterial burden in tissues taken 36 hours post-*Kp* infection. BCKA-treated mice did not show a decline in bacterial colony forming units (CFUs) in spleen, kidney, or lung, suggesting that BCKA treatment was not amplifying the bactericidal capacity of the immune response (**Figure 2E**). To validate these effects, we used the cecal ligation and puncture (CLP) polymicrobial sepsis model. Similar to the *Kp* model, BCKA supplementation rescued survival in a subset of CLP mice without impacting bacterial burden in tissues taken 24 hours post-CLP (**Figure 2F, G**). These results further support the protective effect of BCKA supplementation during sepsis, yet not through augmenting pathogen clearance.

Next, we asked if the protective function of BCKAs is mediated via their catabolism by BCKDC. BT2 is an allosteric inhibitor of branched-chain ketoacid dehydrogenase kinase (BCKDK), which negatively regulates the activity of BCKDC^39^. Thus, BT2 enhances BCKDC activity by inhibiting BCKDK, and was shown to effectively increase BCKDC activity and promote BCKA oxidation in various tissues and disease models such as diet-induced obesity and heart failure^39–43^. When administered alone during Kp sepsis, BT2 did not promote survival compared to the vehicle group (**Figure 2H**). Moreover, BT2 treatment in addition to BCKA supplementation abrogated the protective effect of BCKA supplementation as a monotherapy (**Figure 2H**). This was consistent with increased bacterial load in tissues harvested from BT2-treated mice at endpoint compared to BCKA supplementation (**Figure 2I**). Together, we conclude that the protective effects of BCKA supplementation during sepsis are not mediated by their cellular catabolism.

### BCKAs prevent lipid peroxidation by neutralizing hydrogen peroxide

Our results suggested that BCKAs may be protective in sepsis through a tissue tolerance mechanism and might operate systemically. As lipid peroxidation is one of the prominent pathological features of sepsis^25,34,35^, we assessed the impact of BCKA supplementation on lipid peroxidation in liver tissue by staining 4-hydroxynonenal (4-HNE), a product of lipid peroxidation. Control PBS-treated mice showed high 4-HNE staining particularly at hepatocyte cell junctions 36 hours post *Kp* infection (**Figure 3A**). However, 4-HNE staining was markedly decreased in BCKA-treated mice, suggesting that BCKAs inhibit lipid peroxidation in the liver. To assess whether lipid peroxidation was reduced in bulk liver tissue, we performed a TBARS assay to measure malondialdehyde (MDA), an end product of lipid peroxidation. We found that the liver tissue in BCKA-treated mice during either *Kp*-induced sepsis or CLP sepsis had decreased lipid peroxidation levels (**Figure 3B**).

**Figure 3.**
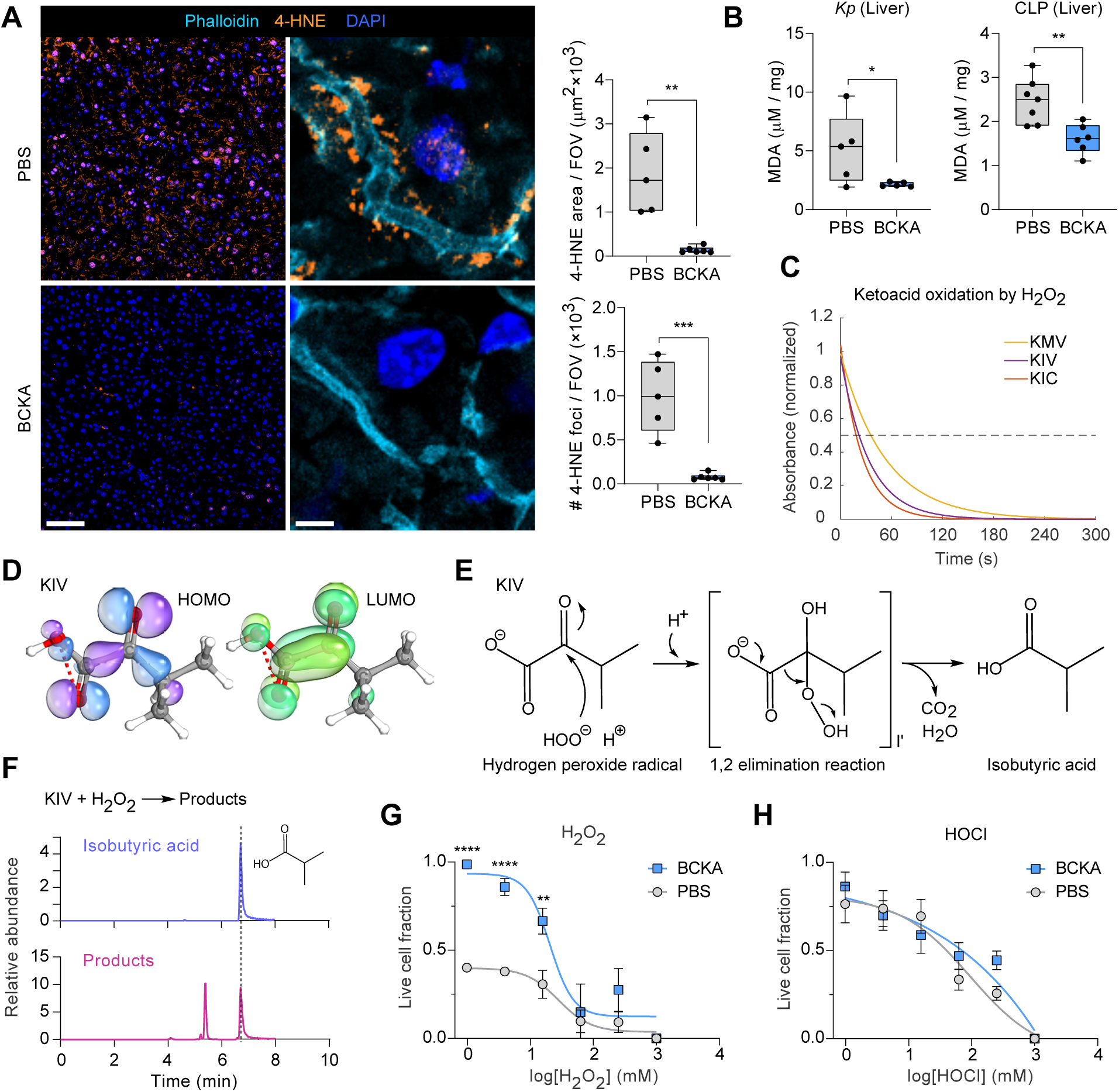
BCKAs prevent lipid peroxidation by neutralizing hydrogen peroxide. (A) Representative immunofluorescence images of liver tissue sections harvested 36 h post *Kp* infection and stained for F-actin (phalloidin) and 4-hydroxynonenal (4-HNE), a product of lipid peroxidation. Scale=50µm (left panels), 5µm (right panels). (*n*=6). (B) Thiobarbituric acid reactive substances (TBARS) assay measuring liver tissue malondialdehyde (MDA), an end product of lipid peroxidation. (C) Ultraviolet–visible spectroscopy (UV-Vis) analysis of BCKA rection with H_2_O_2_. (D) Density functional theory (DFT) analysis of KIV showing the highest occupied molecular orbital (HOMO) and lowest unoccupied molecular orbital (LUMO). (E) Chemical reaction scheme of the putative nucleophilic decarboxylation of KIV by H_2_O_2_. (F) LC-MS analysis of isobutyric acid (top panel) and KIV + H_2_O_2_ reaction products (bottom panel). (G) Cell viability of immortalized bone marrow-derived macrophages (iBMDMs) treated with media containing increasing H_2_O_2_ concentration preincubated with BCKAs or PBS. (H) Cell viability of iBMDMs treated with media containing increasing HOCl concentration preincubated with BCKAs or PBS. Data shown as mean±SEM. *p<0.05; **p<0.01; ***p<0.001; ****p<0.0001.

BCKAs have a unique structure of an α-keto acid, that has the capacity to scavenge hydrogen peroxide (H_2_O_2_)^44^. As the α-keto group is a chromophore that absorbs light at ∼310nm, we monitored the stability of BCKA α-keto group in the presence of peroxide using UV-Vis spectroscopy. All BCKA species had very fast reactions to H_2_O_2_ with t_1/2_ of 30-40 seconds, suggesting direct oxidation by the reaction with peroxide (**Figure 3C**). To quantify how well the BCKAs can be oxidized compared to other antioxidants, we performed density functional theory (DFT) studies using the B3YLP basis set and water as solvent (**Figure 3D**). We modeled KIV and cysteine as the two antioxidants, and we calculated the oxidation potential of KIV to be −1.83 V and cysteine to be −3.28 V. Since KIV’s oxidation potential is much greater than cysteine’s, we would expect it to be better oxidized by peroxide. We then visualized the highest occupied molecular orbital (HOMO) and the lowest unoccupied molecular orbital (LUMO) and found that LUMO overlapped with KIV’s α-carbon, suggesting it may be susceptible to nucleophilic attack. Together with work showing that pyruvate’s α-carbon can undergo a nucleophilic attack by peroxide^45^, we proposed that peroxides can launch a nucleophilic attack and decarboxylate BCKAs, in turn producing short chain fatty acids and CO_2_ (**Figure 3E**). Indeed, LC-MS analysis of the KIV+ H_2_O_2_ reaction products confirmed the production of the short chain fatty acid isobutyric acid—the expected product of KIV decarboxylation (**Figure 3F**).

Because the BCKA decarboxylation reaction is expected to scavenge peroxide, we next tested whether BCKAs can protect cells from peroxide-mediated cytotoxicity by acting as sacrificial ligands. As such, we preincubated BCKAs with increasing doses of H_2_O_2_ or hypochlorous acid (HOCl)—an oxidant that cannot decarboxylate BCKAs. We then added this to media of immortalized bone marrow-derived macrophages (iBMDMs) and measured cell viability using Sytox Green. H_2_O_2_-induced cell death was greatly attenuated following preincubation of H_2_O_2_ with BCKAs (**Figure 3G**). However, BCKA preincubation with HOCl failed to rescue cell viability upon challenge (**Figure 3H**). These results suggest that the protective effect of BCKAs could be attributed to its ability to neutralize peroxide reactivity via a decarboxylation reaction.

### BCKA depletion in human sepsis is associated with poor survival

Low circulating levels of the BCAAs valine, leucine, and isoleucine, have been associated with poor survival and organ failure in sepsis and septic shock patients^46,47^. To establish the relevance of our findings to human sepsis, we analyzed plasma samples collected from 30 patients infected with either *Escherichia coli* or *Kp* during the first 72 hours post admission, as well as 30 age-matched non-septic control patients. All patients were enrolled in the IRB-approved Brigham and Women’s Registry of Critical Illness^48^. LC-MS/MS analysis revealed no remarkable differences in circulating BCAA levels between S and NS patients (**Figure 4A**). However, we found that NS patients had consistently lower levels of BCKAs, particularly KIC and KIV (**Figure 4A**). To determine whether a similar association extends to viral sepsis, we reanalyzed the multi-omics COVID-19 patient dataset published by Su *et al*^49^. Consistent with our findings, severe COVID-19 patients had lower BCKA levels, in particular KIC and KMV, compared to patients with mild or moderate disease (**Figure 4B**). Thus, BCKAs may serve as early prognostic biomarkers in sepsis.

**Figure 4.**
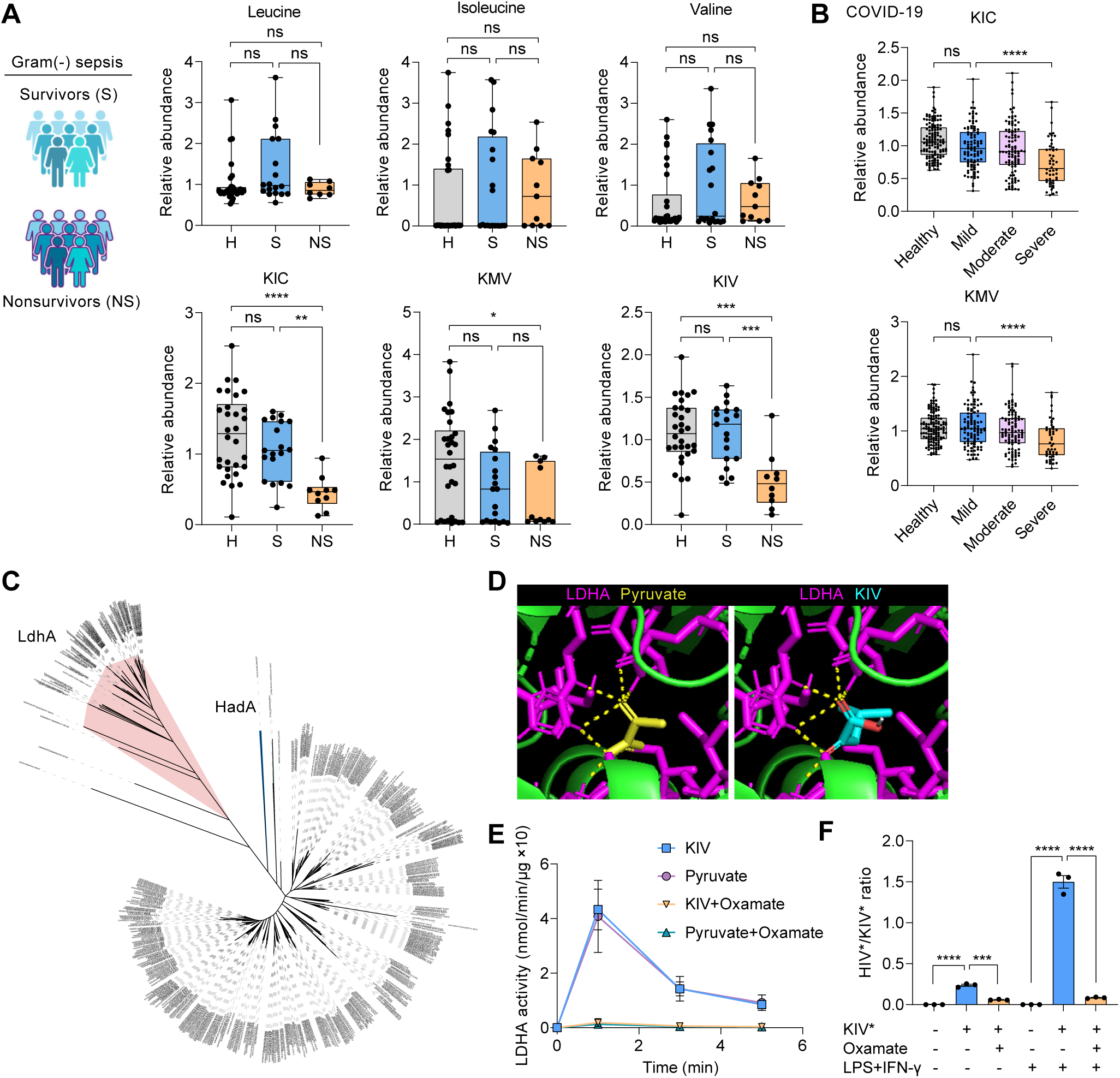
BCKA depletion in human sepsis is associated with poor survival. (A) LC-MS/MS analysis of BCAAs (top panels) or BCKAs (bottom panel) in plasma obtained within 72 h of admission from patients who survived (S) or succumbed to (NS) Gram(-) sepsis, or from healthy non-septic patients (H). (B) LC-MS/MS reanalysis of BCKAs in plasma of COVID-19 patients reported by Su *et al*^49^. (C) Phylogenetic tree of proteins related to the microbial enzyme 2-hydroxyisocaproate dehydrogenase (HadA). LdhA, lactate dehydrogenase A. (D) Model of KIV or pyruvate interaction with human LDHA. Dotted lines indicate putative hydrogen bonding molecular interactions. (E) NADH oxidation to NAD+ by recombinant human LDHA in the presence of pyruvate or KIV and the pyruvate analogue sodium oxamate. (F) LC-MS/MS analysis of the conversion of isotope-labeled KIV (KIV*) to HIV* in iBMDMs culture media. Cells were polarized for 24 h with LPS (50 ng/ml) and interferon (IFN)-γ (20 ng/ml). Data shown as mean±SEM. *p<0.05; **p<0.01; ***p<0.001; ****p<0.0001.

We next asked what may contribute to decreased levels of BCKAs in NS sepsis patients. In bacteria, the enzyme 2-hydroxyisocaproate dehydrogenase (HadA), an oxidoreductase^50^, is responsible for reversibly converting BCKAs to the three branched-chain hydroxyacids (BCHAs) 2-Hydroxy-4-methylpentanoic acid (2-Hydroxyisocaproic acid, HIC), 2-Hydroxy-3-methylpentanoic acid (2-Hydroxy-3-methylvaleric acid, HMV), and 2-Hydroxy-3-methylbutanoic acid (2-Hydroxyisovaleric acid, HIV). To identify mammalian orthologues that may mediate BCKA to BCHA conversion during sepsis, we performed multiple sequence alignment and constructed a phylogenetic tree of proteins related to the microbial HadA. Among 503 related protein sequences, the enzyme lactate dehydrogenase (LdhA) that catalyzes the reversible conversion of pyruvate to lactate was found to be closely related to HadA (**Figure 4C**). Modeling the structural association of KIV or pyruvate with LDHA’s active site highlighted similar stabilizing molecular interactions via hydrogen bonds (**Figure 4D**). To assess whether LDHA could utilize KIV as a substrate similar to pyruvate, we measured the oxidation of the LDHA necessary cofactor NADH by recombinant human LDHA in the presence of either KIV or pyruvate and found similar reaction kinetics (**Figure 4E**). Notably, LDHA activity on both substrates was inhibited by the pyruvate analogue sodium oxamate. We next asked if the cellular conversion of BCKA to BCHA is mediated by LDHA. Incubating iBMDMs with isotope-labeled ^13^C-KIV (KIV*) led to increase in labeled HIV*/KIV* ratio in the culture supernatants, indicating KIV to HIV conversion (**Figure 4F**). However, HIV*/KIV* ratio was reduced in the presence of oxamate. Following LPS and IFN-γ stimulation—which induces glycolytic reprogramming and LDHA in BMDMs^51^—HIV*/KIV* ratio markedly increased, yet not in the presence of oxamate (**Figure 4F**). Taken together, these results suggest that excessive enzymatic depletion of BCKAs may compromise disease tolerance and survival in sepsis.

## DISCUSSION

A multitude of host defenses collapse during sepsis, including immune cell-mediated antimicrobial pathways, and nonimmune mechanisms designed to safeguard tissue integrity in the face of deleterious responses. Antioxidant control is an essential host defense system that enables utilizing free radical-intensive mechanisms of pathogen elimination that would otherwise cause overwhelming damage to the host. Genetic knockout of glutathione peroxidase in myeloid cells underscored the central role of antioxidant systems in preventing sepsis mortality due to increased lipid peroxidation and cell death^52^. Paradoxically, common bactericidal antibiotics increase cellular peroxide production due to mitochondrial oxidative damage^53^. Peroxide is also released by various bacterial species including *S. pneumoniae* and *P. aeruginosa*, leading to cell death, immune dysfunction, and bacterial dissemination^54,55^. It is therefore not surprising that bolstering the host antioxidant capacity in sepsis has become a major therapeutic focus—albeit so far with limited success. For instance, N-acetylcysteine, which replenishes cysteine for glutathione reserves, was ineffective in reducing mortality and complications in sepsis patients and was even harmful by causing cardiovascular depression^37,38^. Similarly, a large study evaluating the therapeutic potential of vitamin C in sepsis showed higher risk of death or persistent organ dysfunction in patients who received intravenous vitamin C compared to those receiving placebo^36^. These failures imply that approaches that bolster endogenous antioxidants may overcome cellular toxicity and enable disease tolerance without compromising early resistance to sepsis. Our data identifies BCKAs as endogenous antioxidants that may act predominantly in the extracellular space, thus mostly preventing lipid peroxidation due to the release of extracellular peroxide.

In support of this notion, there have been attempts to directly neutralize peroxide in the clinic. A clinical trial in cardiac surgery patients used ethyl pyruvate as an antioxidant that undergoes decarboxylation by peroxide, similar to the protective BCKA mechanism we describe^56^. While it was well tolerated, it failed to improve the outcome of the cardiac patients. This is most likely due to steric hindrance introduced by the ethyl ester, preventing the ethyl pyruvate from undergoing the decarboxylation reaction with peroxide. Only intracellularly, the ester is cleaved to form free pyruvate that could undergo decarboxylation. This suggests that in the context of sepsis, where systemic peroxide levels were shown to be elevated both serum and urine^34,57^, neutralizing systemic rather than intracellular peroxide may be more effective in improving survival. A key caveat is that once sufficient damage occurs, antioxidant treatment frequently fails to stop the course of tissue harm as other mechanisms become the dominant drivers of pathology^58,59^. Thus, high preexisting levels of circulating BCKAs may help limit tissue damage early after infection begins to spread. This is reflected by the increased survival of patients with high BCKA levels within 72 hours of admission. BCKAs are mostly stored in skeletal muscle cells and thus decline with age due to a reduction in muscle mass^60^. Lower BCKA stores in elderly patients may thus compromise disease tolerance and contribute to age-related predisposition to sepsis^61^.

Early biomarkers in sepsis patients that predict mortality or organ dysfunction may identify suitable patients for therapeutic intervention^62^. Traditionally, sepsis biomarkers consisted of immune mediators, yet the dynamic nature of the host immune response with oscillating and crossing waves of pro- and anti-inflammatory responses limits the utility of immune biomarker profiles^63,64^. There is now rapidly growing adoption of multi-omics data, including transcriptomics, proteomics, and host and pathogen metagenomics, to identify new biomarkers for sepsis^12,65,66^. Likewise, metabolite profiles have been used to discriminate sepsis from sterile inflammation and predict disease outcomes in sepsis patients^18,19^. Recent studies further highlight the use of machine learning techniques to identify metabolites with prognostic value^67^. A major limitation of sepsis metabolite research is the almost exclusive use of blood samples for analysis, as dictated by patient care considerations. However, tissue metabolite landscapes could provide important insight into disease processes that may be difficult to disentangle solely based on metabolite profiles in circulation. For example, circulating LDHA is commonly used as a sepsis prognostic biomarker associated with poor outcome^68,69^. Our studies interrogating liver metabolite profiles during sepsis uncovered a new functional role for extracellular LDHA in inhibiting the protective antioxidant function of BCKAs. Combining tissue metabolomics and circulating markers of disease could therefore be particularly informative for future discovery of prognostic biomarkers and key molecular pathways in sepsis.

## Acknowledgements

We thank members of the Kanarek and Nowarski laboratories for helpful discussion and critical reading of the manuscript. This work was supported by grants made available to R.N. by the NIH (R35GM133800) and the Gene Lay Institute of Immunology and Inflammation; to N.K. by the Boston Children’s Hospital Pilot Award, and the Smith Family Foundation Excellence Award; to A.M. by the Army Research Office (ARO) in the Department of Defense (DoD) through the National Defense Science & Engineering Graduate (NDSEG) Fellowship Program; and to E.M.W. by The Rothschild Fellowship, Yad-Hanadiv. Measurements in this manuscript were supported by the Neurotechnology Studio at Brigham and Women’s Hospital. N.K. is a Pew Scholar.

## Author contributions

Conceptualization and design of the project – A.M., M.W., E.M.W., N.K., and R.N.; Experimentation and data analysis – A.M., M.W., E.M.W., J.R.B., J.M.R., B.P., R.T.M., N.R., N.K., and R.N.; Provision of key resources – K.V., M.A.P., R.M.B., N.K., and R.N.; Funding and supervision of the project – N.K., and R.N.; Writing the manuscript – A.M., and R.N.; Editing the manuscript – A.M., M.W., E.M.W., N.K., and R.N.

## Methods

### *Kp*-induced sepsis

The wild-type *Klebsiella pneumoniae* subsp. *pneumoniae* (Schroeter) Trevisan (*Kp*) was obtained from ATCC (Cat# 43816). For infection studies, a bacterial culture starter was prepared by placing a stab of bacterial glycerol stock into 3 mL of Nutrient Broth (NB) (BD) for overnight culture (16-18 h at 37°C on a shaker at 200 rpm). The next day, a culture was prepared by putting 200 μL of the culture starter into 20 mL NB and incubating at 37°C and 200 rpm for 2.5 h until the OD600 measured between 0.6-0.8 (logarithmic phase of growth). Bacterial density was estimated using the OD600 measurement, then diluted in PBS to the desired input colony forming unit (CFU) dose (15-100 CFUs in 100 μL PBS per mouse unless otherwise noted). Infection dose was validated by incubating a 100 μL bacterial dilution aliquot on a Nutrient Agar plate (BD) at 37°C for 12 h and counting CFUs. C57BL/6J male and female mice, 7–9 weeks of age, were infected by intraperitoneal injection of 100 μL of *Kp* solution. Rectal temperatures were measured every 4-6 h and body weight was monitored each morning. At any point, mice developing severe hypothermia were sacrificed. For tissue CFU analysis, mouse organs were weighed then homogenized in 1 mL of PBS using the GentleMACS Dissociator (Miltenyi Biotec). The homogenate was serially diluted 1:10 and then 5 μL of each dilution was plated onto an agar plate. After overnight incubation at 37°C, CFUs were enumerated the following day.

### CLP-induced sepsis

C57BL/6J male mice, 7–9 weeks of age, purchased from Charles River, underwent cecal ligation and puncture (CLP) surgery with two-thirds of the cecum ligated and punctured with two 21-gauge holes. In sham experiments, surgery was performed without CLP. The mice received a BCKA cocktail at 1:1:1 molar ratio of KIV, KIC, and KMV (1 mmol/Kg in total) or PBS via daily intraperitoneal administration. Depending on the experiment, the mice were sacrificed at 24 h after CLP for organ CFU analysis or monitored over 4 days to determine survival. In non-survival experiments, organs (liver, spleen, kidney, lung), peritoneal fluid, and blood were harvested for further analyses.

### Metabolomics

Metabolite profiling of mouse liver tissue was performed by the Metabolon global metabolomics platform. Samples were prepared using the automated MicroLab STAR® system from Hamilton Company. Several recovery standards were added prior to the first step in the extraction process for QC purposes. To remove protein, dissociate small molecules bound to protein or trapped in the precipitated protein matrix, and to recover chemically diverse metabolites, proteins were precipitated with methanol under vigorous shaking for 2 min (Glen Mills GenoGrinder 2000) followed by centrifugation. The resulting extract was divided into five fractions: two for analysis by two separate reverse phase (RP)/UPLC-MS/MS methods with positive ion mode electrospray ionization (ESI), one for analysis by RP/UPLC-MS/MS with negative ion mode ESI, one for analysis by HILIC/UPLC-MS/MS with negative ion mode ESI, and one sample was reserved for backup. Samples were placed briefly on a TurboVap® (Zymark) to remove the organic solvent. The sample extracts were stored overnight under nitrogen before preparation for analysis.

#### Ultrahigh Performance Liquid Chromatography-Tandem Mass Spectroscopy (UPLC-MS/MS)

All methods utilized a Waters ACQUITY ultra-performance liquid chromatography (UPLC) and a Thermo Scientific Q-Exactive high resolution/accurate mass spectrometer interfaced with a heated electrospray ionization (HESI-II) source and Orbitrap mass analyzer operated at 35,000 mass resolution. The sample extract was dried then reconstituted in solvents compatible to each of the four methods. Each reconstitution solvent contained a series of standards at fixed concentrations to ensure injection and chromatographic consistency. One aliquot was analyzed using acidic positive ion conditions, chromatographically optimized for more hydrophilic compounds. In this method, the extract was gradient eluted from a C18 column (Waters UPLC BEH C18-2.1x100 mm, 1.7 µm) using water and methanol, containing 0.05% perfluoropentanoic acid (PFPA) and 0.1% formic acid (FA). Another aliquot was also analyzed using acidic positive ion conditions, however it was chromatographically optimized for more hydrophobic compounds. In this method, the extract was gradient eluted from the same afore mentioned C18 column using methanol, acetonitrile, water, 0.05% PFPA and 0.01% FA and was operated at an overall higher organic content. Another aliquot was analyzed using basic negative ion optimized conditions using a separate dedicated C18 column. The basic extracts were gradient eluted from the column using methanol and water, however with 6.5mM Ammonium Bicarbonate at pH 8. The fourth aliquot was analyzed via negative ionization following elution from a HILIC column (Waters UPLC BEH Amide 2.1x150 mm, 1.7 µm) using a gradient consisting of water and acetonitrile with 10mM Ammonium Formate, pH 10.8. The MS analysis alternated between MS and data-dependent MSn scans using dynamic exclusion. The scan range varied slightly between methods but covered 70-1000 m/z. Raw data files are archived and extracted as described below.

#### Data Extraction and Compound Identification

Raw data was extracted, peak-identified and QC processed using Metabolon’s hardware and software. These systems are built on a web-service platform utilizing Microsoft’s .NET technologies, which run on high-performance application servers and fiber-channel storage arrays in clusters to provide active failover and load-balancing. Compounds were identified by comparison to library entries of purified standards or recurrent unknown entities. Metabolon maintains a library based on authenticated standards that contains the retention time/index (RI), mass to charge ratio (m/z), and chromatographic data (including MS/MS spectral data) on all molecules present in the library. Furthermore, biochemical identifications are based on three criteria: retention index within a narrow RI window of the proposed identification, accurate mass match to the library +/- 10 ppm, and the MS/MS forward and reverse scores between the experimental data and authentic standards. The MS/MS scores are based on a comparison of the ions present in the experimental spectrum to the ions present in the library spectrum. While there may be similarities between these molecules based on one of these factors, the use of all three data points can be utilized to distinguish and differentiate biochemicals. More than 3300 commercially available purified standard compounds have been acquired and registered into LIMS for analysis on all platforms for determination of their analytical characteristics. Additional mass spectral entries have been created for structurally unnamed biochemicals, which have been identified by virtue of their recurrent nature (both chromatographic and mass spectral). These compounds have the potential to be identified by future acquisition of a matching purified standard or by classical structural analysis.

### Immortalized bone marrow-derived macrophage (iBMDM) cell death assay

iBMDMs were seeded into a 96 well plate at 50,000 cells/well. Serial dilutions of H_2_O_2_ or HOCl were prepared in normal media or in 10 mM BCKAs. After 1 h incubation at 37°C, the media was then provided to iBMDMs for overnight incubation. Sytox Green staining was performed by staining at 1 μM for 30 min, followed by reading on a plate reader. Cells were lysed using 0.1% Triton-X for 15 min and then another reading was taken.

### Kinetics studies of peroxide reaction

Kinetics was performed by examining the change in absorbance at 310 nm. using a UV-Vis spectrophotometer for 10 minutes with a reading acquired every 10 sec. 900 μL of 100 mM BCKA was added into a plastic microcuvette. The scan was started and then 100 μL of 10M H_2_O_2_ were added into the cuvette such that the H_2_O_2_ had a final concentration of 1M and was in molar excess.

### Measurement of lipid peroxidation using malondialdehyde (MDA) detection kit

The TBARS Assay Kit (TCA method) was used as per manufacturer’s directions (Cayman #700870). Briefly, 15-20 mg pieces of tissue (tissue weights were recorded) were lysed in 250 μL RIPA buffer consisting of 50 mM Tris base (pH = 7.5), 150 mM NaCl, 1 mM EDTA (pH = 8.0), 1% IGEPAL CA-630 (NP-40; Sigma-Aldrich), 0.5% sodium deoxycholate (Sigma-Aldrich) and 0.1% SDS in distilled water with Halt protease inhibitor cocktail (ThermoFisher # 78430) as per the manufacturers’ instructions. Homogenization was performed using a D1000 Handheld Homogenizer (Benchmark Scientific) on ice. Then, the TBARS reagent was added to each sample and incubated at 95°C for 1 h before placing on ice. Samples were centrifuged at 1,600 g for 10 min at 4°C. Supernatant was transferred to 96 well plate and absorbance at 560 nm was read. Absorbance values were normalized to tissue weight.

### Density Functional Theory (DFT) modeling

ORCA was used to perform DFT studies. KIV and cysteine molecules were drawn separately in Avogadro, which then generated the input file for ORCA. First, we performed geometry optimization of the molecules using BP86 as the basis set. After this initial geometry optimization, then the B3YLP basis set was used for geometry optimization calculations. Next, water was specified as the solvent using the CPCMC method in ORCA and geometry optimization and frequency calculations were performed. IboView was used to visualize molecular orbitals. The following equation was used to calculate reduction potential:

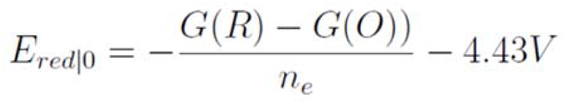

where G(O) is the Gibbs Free Energy of the oxidized species and G(R) is the Gibbs Free Energy of the reduced species. 4.43 V is the standard reduction potential of hydrogen as reference.

### Confocal Microscopy and Immunofluorescence

Fresh tissues were embedded in OCT (Sakura) and frozen on dry ice, then kept in −80°C until sectioning. Tissue sections (8-10 μm) were prepared with a cryostat (Leica) and mounted on SuperFrost Gold slides (ThermoFisher). Sections used for immunofluorescence were fixed with 4% paraformaldehyde (EMS) in PBS for 10 min, permeabilized with Perm/Wash buffer (BD Biosciences) for 10 min, blocked with Serum-Free Protein Blocker (Agilent Dako) for 30 min and incubated with primary antibodies in Perm/Wash buffer for 1h. After three washes with Perm/Wash buffer, sections were stained with secondary antibodies for 1 h, washed 3 times and stained with DAPI (Sigma) for 5 min. Sections were washed with Perm/wash buffer and TrueBlack (Biotium) was added to cover each section for 5 minutes. Sections were washed three times, then mounted with Fluoromount-G Mounting Medium (Southern Biotech) and sealed with nail polish.

## References

1. Singer, M., Deutschman, C.S., Seymour, C.W., Shankar-Hari, M., Annane, D., Bauer, M., Bellomo, R., Bernard, G.R., Chiche, J.-D., and Coopersmith, C.M. (2016). The third international consensus definitions for sepsis and septic shock (Sepsis-3). Jama 315, 801–810.

2. Rhee, C., Jones, T.M., Hamad, Y., Pande, A., Varon, J., O’Brien, C., Anderson, D.J., Warren, D.K., Dantes, R.B., and Epstein, L. (2019). Prevalence, underlying causes, and preventability of sepsis-associated mortality in US acute care hospitals. JAMA network open 2, e187571–e187571.

3. Rudd, K.E., Johnson, S.C., Agesa, K.M., Shackelford, K.A., Tsoi, D., Kievlan, D.R., Colombara, D.V., Ikuta, K.S., Kissoon, N., and Finfer, S. (2020). Global, regional, and national sepsis incidence and mortality, 1990–2017: analysis for the Global Burden of Disease Study. The Lancet 395, 200–211.

4. Medzhitov, R., Schneider, D.S., and Soares, M.P. (2012). Disease tolerance as a defense strategy. Science 335, 936–941.

5. van der Poll, T., Shankar-Hari, M., and Wiersinga, W.J. (2021). The immunology of sepsis. Immunity 54, 2450–2464.

6. van der Poll, T., van de Veerdonk, F.L., Scicluna, B.P., and Netea, M.G. (2017). The immunopathology of sepsis and potential therapeutic targets. Nat Rev Immunol 17, 407–420. 10.1038/nri.2017.36.

7. Burnham, K.L., Davenport, E.E., Radhakrishnan, J., Humburg, P., Gordon, A.C., Hutton, P., Svoren-Jabalera, E., Garrard, C., Hill, A.V., and Hinds, C.J. (2017). Shared and distinct aspects of the sepsis transcriptomic response to fecal peritonitis and pneumonia. American journal of respiratory and critical care medicine 196, 328–339.

8. Willmann, K., and Moita, L.F. (2024). Physiologic disruption and metabolic reprogramming in infection and sepsis. Cell metabolism 36, 927–946. 10.1016/j.cmet.2024.02.013.

9. Wang, A., Luan, H.H., and Medzhitov, R. (2019). An evolutionary perspective on immunometabolism. Science 363, eaar3932.

10. Cheng, S.C., Scicluna, B.P., Arts, R.J., Gresnigt, M.S., Lachmandas, E., Giamarellos-Bourboulis, E.J., Kox, M., Manjeri, G.R., Wagenaars, J.A., Cremer, O.L., et al. (2016). Broad defects in the energy metabolism of leukocytes underlie immunoparalysis in sepsis. Nature immunology 17, 406–413. 10.1038/ni.3398.

11. Weis, S., Carlos, A.R., Moita, M.R., Singh, S., Blankenhaus, B., Cardoso, S., Larsen, R., Rebelo, S., Schäuble, S., Del Barrio, L., et al. (2017). Metabolic Adaptation Establishes Disease Tolerance to Sepsis. Cell 169, 1263–1275.e1214. 10.1016/j.cell.2017.05.031.

12. Reyes, M., Filbin, M.R., Bhattacharyya, R.P., Billman, K., Eisenhaure, T., Hung, D.T., Levy, B.D., Baron, R.M., Blainey, P.C., and Goldberg, M.B. (2020). An immune-cell signature of bacterial sepsis. Nature medicine 26, 333–340.

13. Paumelle, R., Haas, J.T., Hennuyer, N., Baugé, E., Deleye, Y., Mesotten, D., Langouche, L., Vanhoutte, J., Cudejko, C., Wouters, K., et al. (2019). Hepatic PPARα is critical in the metabolic adaptation to sepsis. J Hepatol 70, 963–973. 10.1016/j.jhep.2018.12.037.

14. Wang, A., Huen, S.C., Luan, H.H., Yu, S., Zhang, C., Gallezot, J.-D., Booth, C.J., and Medzhitov, R. (2016). Opposing Effects of Fasting Metabolism on Tissue Tolerance in Bacterial and Viral Inflammation. Cell 166, 1512–1525.e1512. 10.1016/j.cell.2016.07.026.

15. Huen, S.C. (2021). Metabolism as Disease Tolerance: Implications for Sepsis-Associated Acute Kidney Injury. Nephron 146, 291–294. 10.1159/000516877.

16. Wasyluk, W., and Zwolak, A. (2021). Metabolic Alterations in Sepsis. J Clin Med 10. 10.3390/jcm10112412.

17. Vandewalle, J., and Libert, C. (2022). Sepsis: a failing starvation response. Trends Endocrinol Metab 33, 292–304. 10.1016/j.tem.2022.01.006.

18. Ferrario, M., Cambiaghi, A., Brunelli, L., Giordano, S., Caironi, P., Guatteri, L., Raimondi, F., Gattinoni, L., Latini, R., and Masson, S. (2016). Mortality prediction in patients with severe septic shock: a pilot study using a target metabolomics approach. Scientific reports 6, 20391.

19. Langley, R.J., Tsalik, E.L., Velkinburgh, J.C.v., Glickman, S.W., Rice, B.J., Wang, C., Chen, B., Carin, L., Suarez, A., Mohney, R.P., et al. (2013). An Integrated Clinico-Metabolomic Model Improves Prediction of Death in Sepsis. Science Translational Medicine 5, 195ra195–195ra195. doi:10.1126/scitranslmed.3005893.

20. Beyer, D., Hoff, J., Sommerfeld, O., Zipprich, A., Gaßler, N., and Press, A.T. (2022). The liver in sepsis: molecular mechanism of liver failure and their potential for clinical translation. Molecular Medicine 28, 84.

21. Yan, J., Li, S., and Li, S. (2014). The role of the liver in sepsis. International reviews of immunology 33, 498–510.

22. Guo, Y., Guo, W., Chen, H., Sun, J., and Yin, Y. (2025). Mechanisms of sepsis-induced acute liver injury: a comprehensive review. Front Cell Infect Microbiol 15, 1504223. 10.3389/fcimb.2025.1504223.

23. Murphy, M.P., Bayir, H., Belousov, V., Chang, C.J., Davies, K.J.A., Davies, M.J., Dick, T.P., Finkel, T., Forman, H.J., Janssen-Heininger, Y., et al. (2022). Guidelines for measuring reactive oxygen species and oxidative damage in cells and in vivo. Nature Metabolism 4, 651–662. 10.1038/s42255-022-00591-z.

24. Rowe, S.E., Wagner, N.J., Li, L., Beam, J.E., Wilkinson, A.D., Radlinski, L.C., Zhang, Q., Miao, E.A., and Conlon, B.P. (2020). Reactive oxygen species induce antibiotic tolerance during systemic Staphylococcus aureus infection. Nature Microbiology 5, 282–290. 10.1038/s41564-019-0627-y.

25. Cowley, H.C., Bacon, P.J., Goode, H.F., Webster, N.R., Jones, J.G., and Menon, D.K. (1996). Plasma antioxidant potential in severe sepsis: a comparison of survivors and nonsurvivors. Critical care medicine 24, 1179–1183.

26. Sahoo, D.K., Wong, D., Patani, A., Paital, B., Yadav, V.K., Patel, A., and Jergens, A.E. (2024). Exploring the role of antioxidants in sepsis-associated oxidative stress: a comprehensive review. Frontiers in Cellular and Infection Microbiology 14, 1348713.

27. Hoetzenecker, W., Echtenacher, B., Guenova, E., Hoetzenecker, K., Woelbing, F., Brück, J., Teske, A., Valtcheva, N., Fuchs, K., and Kneilling, M. (2012). ROS-induced ATF3 causes susceptibility to secondary infections during sepsis-associated immunosuppression. Nature medicine 18, 128–134.

28. Ayala, J.C., Grismaldo, A., Sequeda-Castañeda, L.G., Aristizábal-Pachón, A.F., and Morales, L. (2021). Oxidative Stress in ICU Patients: ROS as Mortality Long-Term Predictor. Antioxidants 10, 1912.

29. Imlay, J.A. (2013). The molecular mechanisms and physiological consequences of oxidative stress: lessons from a model bacterium. Nature Reviews Microbiology 11, 443–454.

30. Biolo, G., Antonione, R., and De Cicco, M. (2007). Glutathione metabolism in sepsis. Critical care medicine 35, S591–S595.

31. Tandon, R., and Tandon, A. (2024). Unraveling the Multifaceted Role of Glutathione in Sepsis: A Comprehensive Review. Cureus 16, e56896. 10.7759/cureus.56896.

32. Callahan, L.A., and Supinski, G.S. (2009). Sepsis-induced myopathy. Critical care medicine 37, S354–367. 10.1097/CCM.0b013e3181b6e439.

33. Pravda, J. (2021). Sepsis: Evidence-based pathogenesis and treatment. World J Crit Care Med 10, 66–80. 10.5492/wjccm.v10.i4.66.

34. Ogilvie, A.C., Groeneveld, A.B., Straub, J.P., and Thijs, L.G. (1991). Plasma lipid peroxides and antioxidants in human septic shock. Intensive care medicine 17, 40–44. 10.1007/bf01708408.

35. Goode, H.F., Cowley, H.C., Walker, B.E., Howdle, P.D., and Webster, N.R. (1995). Decreased antioxidant status and increased lipid peroxidation in patients with septic shock and secondary organ dysfunction. Critical care medicine 23, 646–651. 10.1097/00003246-199504000-00011.

36. Lamontagne, F., Masse, M.-H., Menard, J., Sprague, S., Pinto, R., Heyland, D.K., Cook, D.J., Battista, M.-C., Day, A.G., Guyatt, G.H., et al. (2022). Intravenous Vitamin C in Adults with Sepsis in the Intensive Care Unit. New England Journal of Medicine 386, 2387–2398. doi:10.1056/NEJMoa2200644.

37. Szakmany, T., Hauser, B., and Radermacher, P. (2012). N-acetylcysteine for sepsis and systemic inflammatory response in adults. Cochrane Database Syst Rev 2012, CD006616. 10.1002/14651858.CD006616.pub2.

38. Spapen, H.D., Diltoer, M.W., Nguyen, D.N., Hendrickx, I., and Huyghens, L.P. (2005). Effects of N-acetylcysteine on microalbuminuria and organ failure in acute severe sepsis: results of a pilot study. Chest 127, 1413–1419. 10.1378/chest.127.4.1413.

39. Tso, S.C., Gui, W.J., Wu, C.Y., Chuang, J.L., Qi, X., Skvora, K.J., Dork, K., Wallace, A.L., Morlock, L.K., Lee, B.H., et al. (2014). Benzothiophene carboxylate derivatives as novel allosteric inhibitors of branched-chain α-ketoacid dehydrogenase kinase. The Journal of biological chemistry 289, 20583–20593. 10.1074/jbc.M114.569251.

40. Bollinger, E., Peloquin, M., Libera, J., Albuquerque, B., Pashos, E., Shipstone, A., Hadjipanayis, A., Sun, Z., Xing, G., Clasquin, M., et al. (2022). BDK inhibition acts as a catabolic switch to mimic fasting and improve metabolism in mice. Molecular Metabolism 66, 101611. 10.1016/j.molmet.2022.101611.

41. Wang, W., Zhang, F., Xia, Y., Zhao, S., Yan, W., Wang, H., Lee, Y., Li, C., Zhang, L., Lian, K., et al. (2016). Defective branched chain amino acid catabolism contributes to cardiac dysfunction and remodeling following myocardial infarction. Am J Physiol Heart Circ Physiol 311, H1160–H1169. 10.1152/ajpheart.00114.2016.

42. Zhou, M., Shao, J., Wu, C.Y., Shu, L., Dong, W., Liu, Y., Chen, M., Wynn, R.M., Wang, J., Wang, J., et al. (2019). Targeting BCAA Catabolism to Treat Obesity-Associated Insulin Resistance. Diabetes 68, 1730–1746. 10.2337/db18-0927.

43. Chen, M., Gao, C., Yu, J., Ren, S., Wang, M., Wynn, R.M., Chuang, D.T., Wang, Y., and Sun, H. (2019). Therapeutic Effect of Targeting Branched-Chain Amino Acid Catabolic Flux in Pressure-Overload Induced Heart Failure. J Am Heart Assoc 8, e011625. 10.1161/jaha.118.011625.

44. Nath, K.A., Ngo, E.O., Hebbel, R.P., Croatt, A.J., Zhou, B., and Nutter, L.M. (1995). alpha-Ketoacids scavenge H2O2 in vitro and in vivo and reduce menadione-induced DNA injury and cytotoxicity. The American journal of physiology 268, C227–236. 10.1152/ajpcell.1995.268.1.C227.

45. Lopalco, A., Dalwadi, G., Niu, S., Schowen, R.L., Douglas, J., and Stella, V.J. (2016). Mechanism of Decarboxylation of Pyruvic Acid in the Presence of Hydrogen Peroxide. J Pharm Sci 105, 705–713. 10.1002/jps.24653.

46. Reisinger, A.C., Posch, F., Hackl, G., Marsche, G., Sourij, H., Bourgeois, B., Eller, K., Madl, T., and Eller, P. (2021). Branched-Chain Amino Acids Can Predict Mortality in ICU Sepsis Patients. Nutrients 13. 10.3390/nu13093106.

47. Puskarich, M.A., McHugh, C., Flott, T.L., Karnovsky, A., Jones, A.E., and Stringer, K.A. (2021). Serum Levels of Branched Chain Amino Acids Predict Duration of Cardiovascular Organ Failure in Septic Shock. Shock 56.

48. Dolinay, T., Kim, Y.S., Howrylak, J., Hunninghake, G.M., An, C.H., Fredenburgh, L., Massaro, A.F., Rogers, A., Gazourian, L., Nakahira, K., et al. (2012). Inflammasome-regulated cytokines are critical mediators of acute lung injury. American journal of respiratory and critical care medicine 185, 1225–1234. 10.1164/rccm.201201-0003OC.

49. Su, Y., Chen, D., Yuan, D., Lausted, C., Choi, J., Dai, C.L., Voillet, V., Duvvuri, V.R., Scherler, K., Troisch, P., et al. (2020). Multi-Omics Resolves a Sharp Disease-State Shift between Mild and Moderate COVID-19. Cell 183, 1479–1495 e1420. 10.1016/j.cell.2020.10.037.

50. Niefind, K., Hecht, H.J., and Schomburg, D. (1995). Crystal structure of L-2-hydroxyisocaproate dehydrogenase from Lactobacillus confusus at 2.2 A resolution. An example of strong asymmetry between subunits. J Mol Biol 251, 256–281. 10.1006/jmbi.1995.0433.

51. Mertens, R.T., Misra, A., Xiao, P., Baek, S., Rone, J.M., Mangani, D., Sivanathan, K.N., Arojojoye, A.S., Awuah, S.G., Lee, I., et al. (2024). A metabolic switch orchestrated by IL-18 and the cyclic dinucleotide cGAMP programs intestinal tolerance. Immunity 57, 2077–2094 e2012. 10.1016/j.immuni.2024.06.001.

52. Kang, R., Zeng, L., Zhu, S., Xie, Y., Liu, J., Wen, Q., Cao, L., Xie, M., Ran, Q., and Kroemer, G. (2018). Lipid peroxidation drives gasdermin D-mediated pyroptosis in lethal polymicrobial sepsis. Cell host & microbe 24, 97–108. e104.

53. Kalghatgi, S., Spina, C.S., Costello, J.C., Liesa, M., Morones-Ramirez, J.R., Slomovic, S., Molina, A., Shirihai, O.S., and Collins, J.J. (2013). Bactericidal antibiotics induce mitochondrial dysfunction and oxidative damage in mammalian cells. Science translational medicine 5, 192ra185–192ra185.

54. Erttmann, S.F., and Gekara, N.O. (2019). Hydrogen peroxide release by bacteria suppresses inflammasome-dependent innate immunity. Nature Communications 10, 3493.

55. Rai, P., Parrish, M., Tay, I.J.J., Li, N., Ackerman, S., He, F., Kwang, J., Chow, V.T., and Engelward, B.P. (2015). Streptococcus pneumoniae secretes hydrogen peroxide leading to DNA damage and apoptosis in lung cells. Proceedings of the National Academy of Sciences 112, E3421–E3430.

56. Yang, R., Zhu, S., and Tonnessen, T.I. (2016). Ethyl pyruvate is a novel anti-inflammatory agent to treat multiple inflammatory organ injuries. J Inflamm (Lond) 13, 37. 10.1186/s12950-016-0144-1.

57. Lipcsey, M., Bergquist, M., Sirén, R., Larsson, A., Huss, F., Pravda, J., Furebring, M., Sjölin, J., and Janols, H. (2022). Urine Hydrogen Peroxide Levels and Their Relation to Outcome in Patients with Sepsis, Septic Shock, and Major Burn Injury. Biomedicines 10. 10.3390/biomedicines10040848.

58. Forman, H.J., and Zhang, H. (2021). Targeting oxidative stress in disease: promise and limitations of antioxidant therapy. Nature Reviews Drug Discovery 20, 689–709.

59. Jena, A.B., Samal, R.R., Bhol, N.K., and Duttaroy, A.K. (2023). Cellular Red-Ox system in health and disease: The latest update. Biomedicine & Pharmacotherapy 162, 114606. 10.1016/j.biopha.2023.114606.

60. Mann, G., Mora, S., Madu, G., and Adegoke, O.A.J. (2021). Branched-chain Amino Acids: Catabolism in Skeletal Muscle and Implications for Muscle and Whole-body Metabolism. Front Physiol 12, 702826. 10.3389/fphys.2021.702826.

61. Meyer Nuala, J., and Prescott Hallie, C. (2024). Sepsis and Septic Shock. New England Journal of Medicine 391, 2133–2146. 10.1056/NEJMra2403213.

62. Barichello, T., Generoso, J.S., Singer, M., and Dal-Pizzol, F. (2022). Biomarkers for sepsis: more than just fever and leukocytosis—a narrative review. Critical Care 26, 14. 10.1186/s13054-021-03862-5.

63. Llitjos, J.-F., Carrol, E.D., Osuchowski, M.F., Bonneville, M., Scicluna, B.P., Payen, D., Randolph, A.G., Witte, S., Rodriguez-Manzano, J., François, B., and on behalf of the Sepsis biomarker workshop, g. (2024). Enhancing sepsis biomarker development: key considerations from public and private perspectives. Critical Care 28, 238. 10.1186/s13054-024-05032-9.

64. Hotchkiss, R.S., Monneret, G., and Payen, D. (2013). Sepsis-induced immunosuppression: from cellular dysfunctions to immunotherapy. Nature Reviews Immunology 13, 862–874.

65. Sweeney, T.E., Shidham, A., Wong, H.R., and Khatri, P. (2015). A comprehensive time-course–based multicohort analysis of sepsis and sterile inflammation reveals a robust diagnostic gene set. Science translational medicine 7, 287ra271–287ra271.

66. Kalantar, K.L., Neyton, L., Abdelghany, M., Mick, E., Jauregui, A., Caldera, S., Serpa, P.H., Ghale, R., Albright, J., and Sarma, A. (2022). Integrated host-microbe plasma metagenomics for sepsis diagnosis in a prospective cohort of critically ill adults. Nature microbiology 7, 1805–1816.

67. Komorowski, M., Green, A., Tatham, K.C., Seymour, C., and Antcliffe, D. (2022). Sepsis biomarkers and diagnostic tools with a focus on machine learning. EBioMedicine 86. 10.1016/j.ebiom.2022.104394.

68. Lu, J., Wei, Z., Jiang, H., Cheng, L., Chen, Q., Chen, M., Yan, J., and Sun, Z. (2018). Lactate dehydrogenase is associated with 28-day mortality in patients with sepsis: a retrospective observational study. Journal of Surgical Research 228, 314–321. 10.1016/j.jss.2018.03.035.

69. Zein, J.G., Lee, G.L., Tawk, M., Dabaja, M., and Kinasewitz, G.T. (2004). Prognostic Significance of Elevated Serum Lactate Dehydrogenase (LDH) in Patients with Severe Sepsis. CHEST 126, 873S. 10.1378/chest.126.4_MeetingAbstracts.873S.

